# Tricuspid valve leaflet remodeling in sheep with biventricular heart failure: A comparison between leaflets

**DOI:** 10.1101/2024.09.16.613284

**Authors:** Colton J. Kostelnik, William D. Meador, Chien-Yu Lin, Mrudang Mathur, Marcin Malinowski, Tomasz Jazwiec, Zuzanna Malinowska, Magda L. Piekarska, Boguslaw Gaweda, Tomasz A. Timek, Manuel K. Rausch

**Author notes:** Corresponding Author: Manuel K. Rausch, 2501 Speedway, EER Building, Room 7.620. Colton Kostelnik and William Meador are co-first authors.

## Abstract

Tricuspid valve leaflets are dynamic tissues that can respond to altered biomechanical and hemodynamic loads. Each leaflet has unique structural and mechanical properties, leading to differential in vivo strains. We hypothesized that these intrinsic differences drive heterogeneous, disease-induced remodeling between the leaflets. Although we previously reported significant remodeling changes in the anterior leaflet, the responses among the other two leaflets have not been reported. Using a sheep model of biventricular heart failure, we compared the remodeling responses between all tricuspid leaflets. Our results show that the anterior leaflet underwent the most significant remodeling, while the septal and posterior leaflets exhibited similar but less pronounced changes. We found several between-leaflet differences in key structural and mechanical metrics that have been shown to contribute to valvular dysfunction. These findings underscore the need to consider leaflet-specific remodeling to fully understand tricuspid valve dysfunction and to develop targeted therapies for its treatment and more accurate computational models.

**STATEMENT OF SIGNIFICANCE:** Our study is significant as it advances our understanding of tricuspid valve remodeling by providing a comprehensive analysis of all three leaflets in a sheep model of biventricular heart failure. Unlike prior works that focused primarily on the anterior leaflet or generalized leaflet changes, we integrated morphological, histological, immunohistochemistry, biaxial mechanical testing, and two-photon microscopy to quantify differences between all three tricuspid valve leaflets (anterior, posterior, and septal) across multiple functional scales. This comprehensive approach highlights the unique remodeling response of each leaflet. Our findings offer critical insights for developing targeted therapeutic strategies and improving computational models of disease progression.

**GRAPHICAL ABSTRACT:** 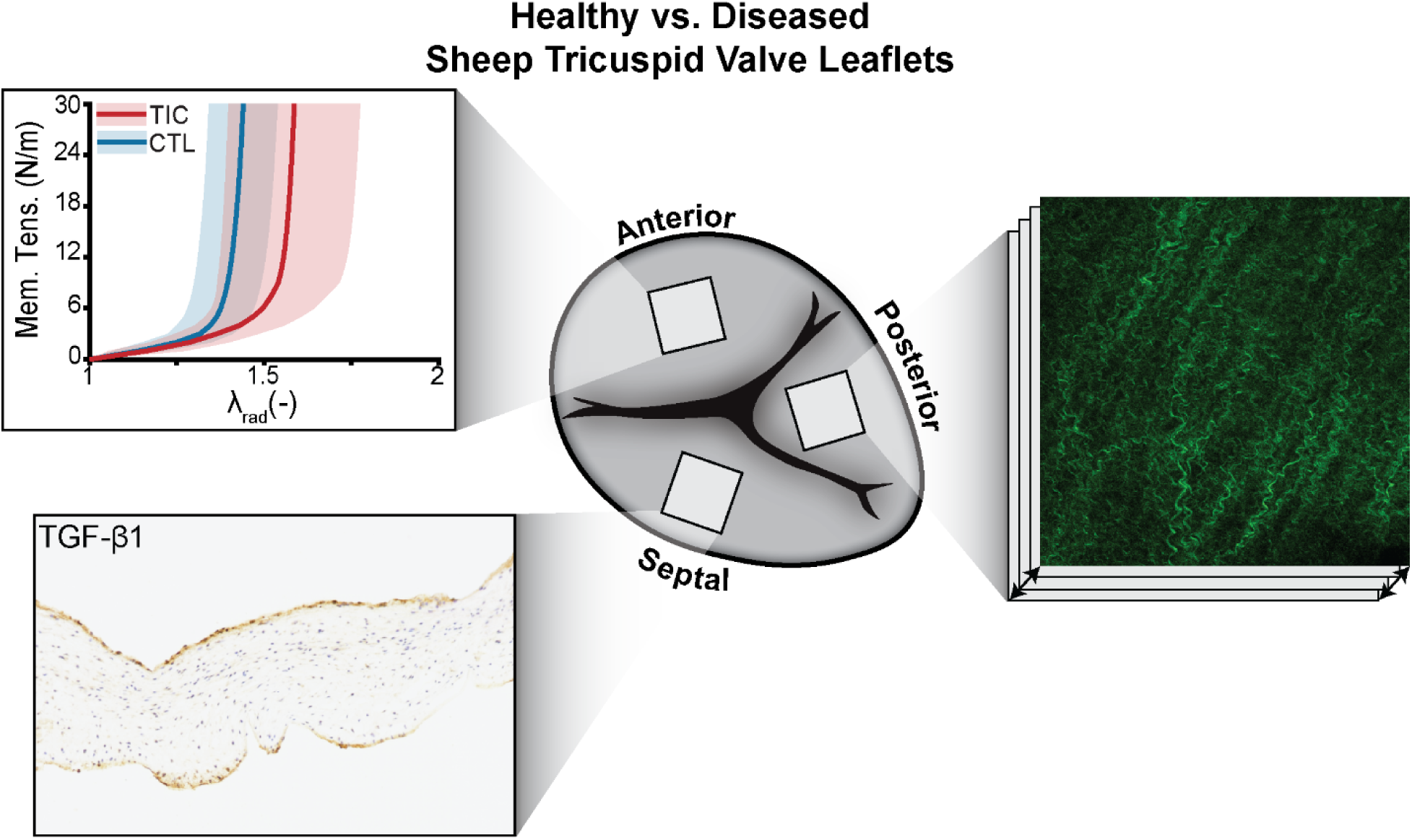

## 1. INTRODUCTION

Tricuspid regurgitation (TR) is a highly prevalent echocardiographic finding detected in 82-86% of people [1]. Approximately 1.6 million Americans suffer from moderate to severe TR, which, if left untreated, can lead to multiple cardiovascular complications [2]. More than 90% of patients with severe TR exhibit a "functional" etiology, wherein the valve dysfunction is secondary to other cardiac pathologies such as left-sided heart disease or pulmonary hypertension [3–5]. The valve itself has historically been considered structurally and mechanically intact, and this assumption has guided a conservative treatment approach that focuses on managing the primary condition to resolve any TR [6,7]. However, early surgical repair of the tricuspid valve during mitral or aortic valve surgeries has shown promise in improving patient outcomes and diminishing cardiac-related mortality [8,9]. Despite this evolution of TR management, it remains undertreated and poorly understood.

Heart valve leaflets were once considered "passive flaps" that merely coordinated unidirectional blood flow between the heart chambers akin to a check valve. However, we now know that heart valve leaflets are highly organized tissues with multiple cell types and intricate extracellular architectures that actively remodel by contracting and reshaping to optimize coaptation [10–12]. In diseased patients, leaflet remodeling induces a fibrotic response (i.e., maladapt), which impedes proper kinematics and coaptation [13]. Animal studies of heart failure and chronic ischemia have shown that maladaptive changes in the mitral valve leaflets manifest by increased size, thickness, and stiffness [13–16]. We have demonstrated in two animal models that similar remodeling patterns exist in the tricuspid valve leaflets (e.g., increases in area, thickness, and stiffness) in response to altered biomechanical and hemodynamic loads [17,18]. These remodeling-induced leaflet changes affect leaflet stresses and leaflet contact area (i.e., coaptation area), indicating that valve function is sensitive to leaflet maladaptation [19].

Moreover, we have previously demonstrated that the tricuspid leaflets in healthy sheep exhibit distinct, leaflet-specific properties, including unique morphology, mechanical properties, and in vivo mechanics [20,21]. In our prior study of tricuspid valve maladaptation, we found that the anterior tricuspid leaflet exhibited significant remodeling changes on all functional scales in sheep with biventricular heart failure after 2-3 weeks of tachycardia-induced cardiomyopathy (TIC) [17]. Building on this work, here we aim to quantify the remodeling changes in the septal and posterior tricuspid leaflets from the same sheep and compare the remodeling patterns across all three leaflets. We hypothesize that the leaflet-specific mechanics drive heterogeneous remodeling responses between the tricuspid leaflets. Specifically, we hypothesize that the TIC anterior leaflets exhibit the most significant remodeling changes relative to its control due to its larger in vivo area strains, while the TIC septal and posterior leaflets remodel to a lesser extent [20]. This distinction of differential remodeling may be vital in uncovering potential therapeutic targets tailored to specific leaflets and informing computational models of disease progression.

## 2. MATERIALS & METHODS

### 2.1 Animal Model, Medication, and Procedures

This research complied in accordance with the Principles of Laboratory Animal Care, formulated by the National Society for Medical Research, and adhered to protocols outlined in the Guide for Care and Use of Laboratory Animals, published by the National Institutes of Health. Additionally, this protocol was developed, reviewed and performed in accordance with the approval of a local Institutional Animal Care and Use Committee at Corewell Health. The specific approval numbers are 2017-357, 2018-402, 2018427, and 2018-439.

The TIC sheep model used in this study has been previously described in detail and demonstrated as a reliable and repeatable model of biventricular heart failure in sheep [22]. Detailed descriptions of the animal model, medications, and procedures are available in our prior work [17]. We randomly assigned adult male Dorsett sheep to either the control (CTL) (n = 17, 59.9 ± 4.6 kg) or TIC (n = 33, 60.1 ± 5.3 kg) groups. Before initiating pacing, we monitored baseline ventricular function and valvular competence in all animals using epicardial echocardiography. The TIC sheep were then subject to a 180-260 beats per minute pacing protocol for 19 ± 6 days. Following the pacing period, we performed the terminal procedure on all TIC sheep and recorded hemodynamic and echocardiographic data over three cardiac cycles. In CTL sheep, we performed the terminal procedure following baseline epicardial echocardiography. We reported all hemodynamic and echocardiographic data in Table 1 of our previous study [17]. After completing the terminal procedures, we isolated the tricuspid valve from each animal for further analysis.

**Table 1:**
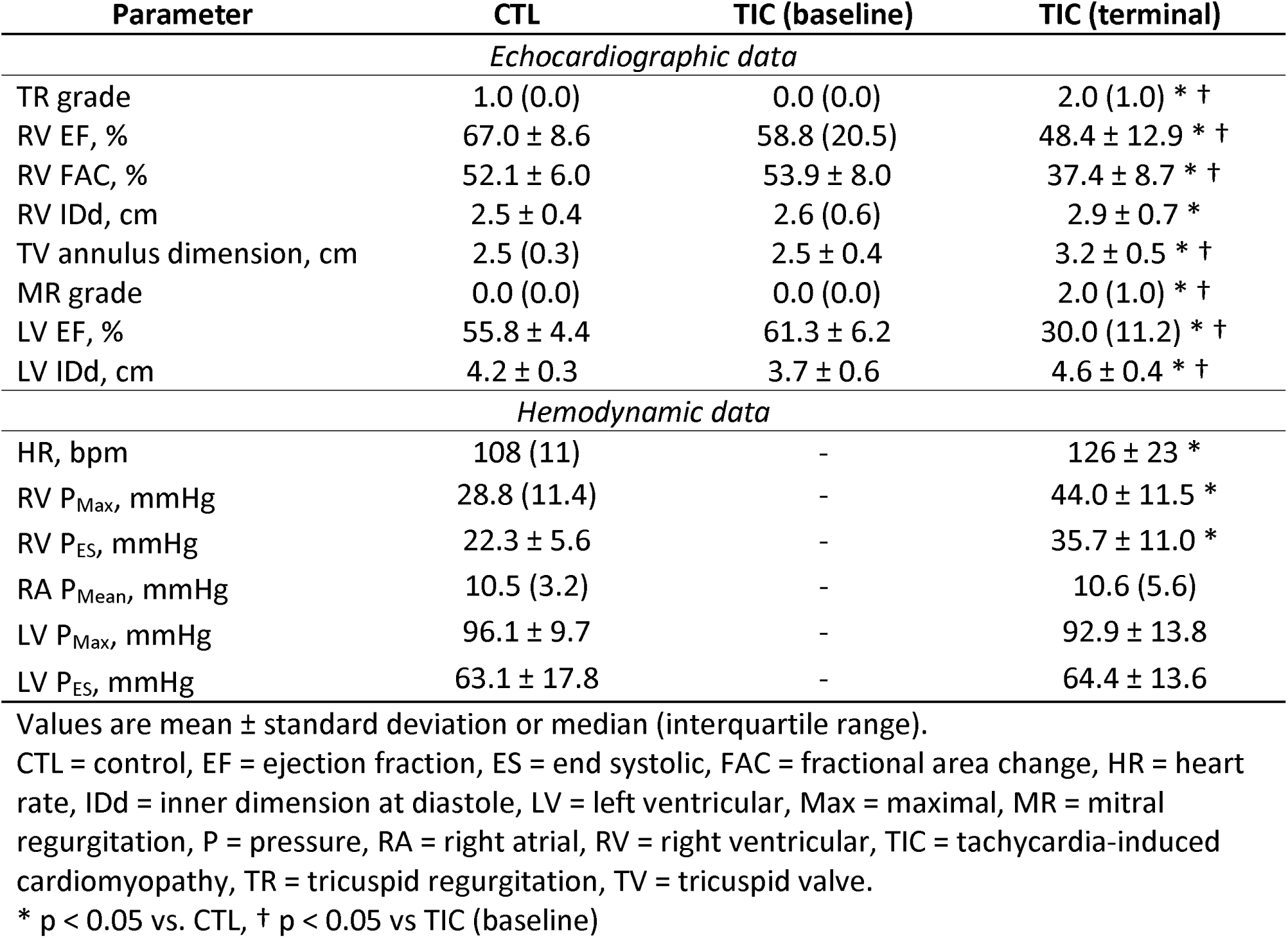
Echocardiographic and hemodynamic data of animal model.

For disclosure, we previously published data on the anterior leaflet morphology, thickness, biaxial mechanics, histology, collagen content, and two-photon microscopy from the sheep included in this study [17]. We include these data here again with expressed permission from the editorial office.

### 2.2 Morphology and Storage

We isolated the tricuspid valve leaflet complex from each heart and separated the leaflets at the posterior-septal commissure. We then floated each valve on 1x PBS and imaged them on a calibrated grid to compare morphological measures such as leaflet area between CTL and TIC animals (Figure 1a). The leaflets were then separated and cryopreserved in DMEM (VWR L0101-0500, Radnor, PA, USA) containing 10% DMSO (VWR BDH1115-1LP) supplemented with a protease inhibitor (ThermoFisher, A32953, Weltham, MA, USA) and stored at -80°C until tested. Cryogenic studies have shown that the mechanical properties of soft tissues were minimally impacted after storage at temperatures below 0°C [23,24].

**Figure 1:**
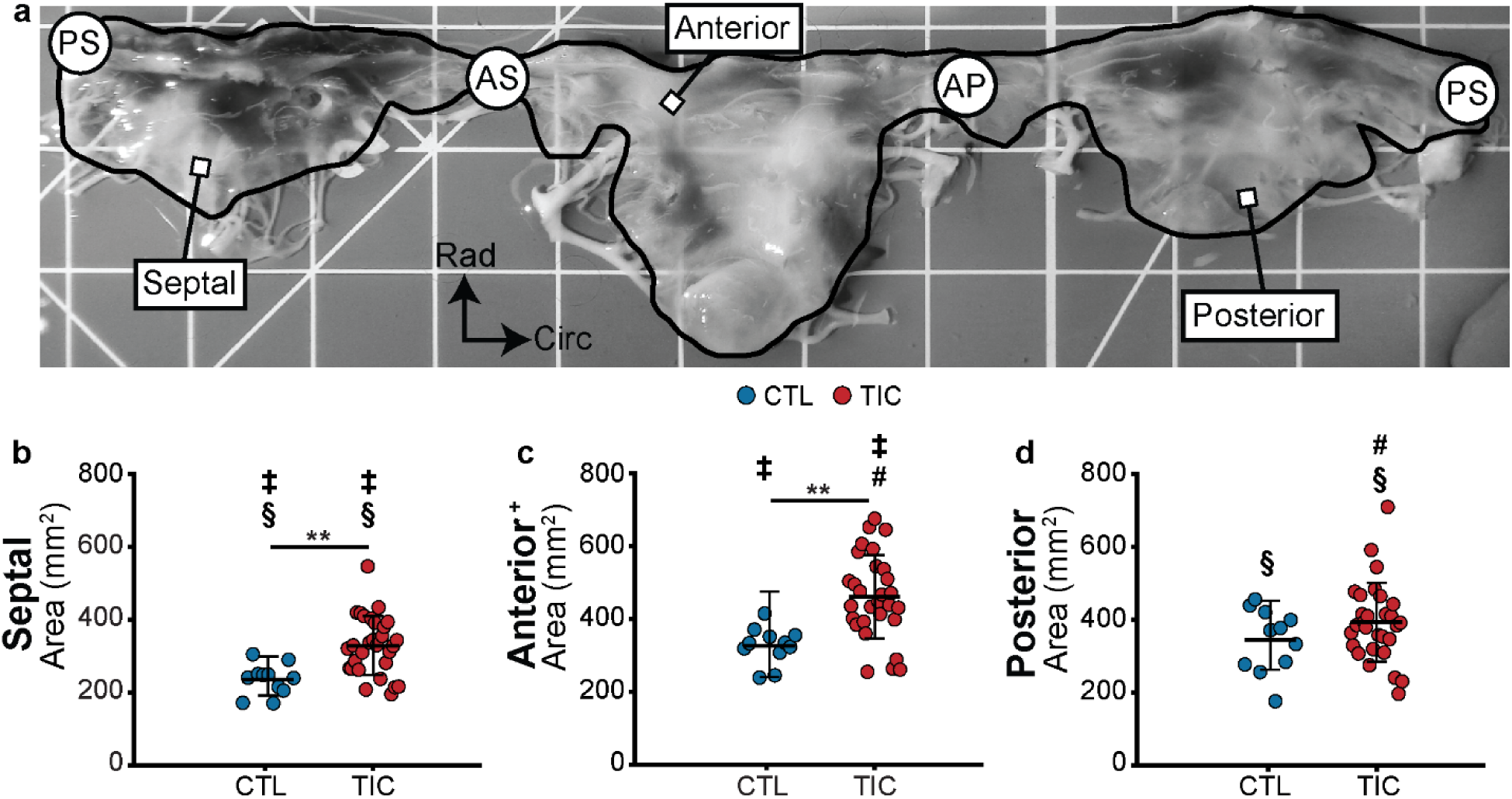
Tricuspid septal and anterior leaflet area increases in sheep with tachycardia-induced cardiomyopathy (TIC). (a) Sheep tricuspid valve image on a calibrated grid (1 cm scale). Measured areas (black line) of septal, anterior, and posterior leaflets are shown. Leaflet areas were measured between commissures: posterior-septal (PS) to antero-septal (AS) for septal, AS to antero-posterior (AP) for anterior, and AP to PS for posterior. (b-d) Leaflet area comparisons between control (CTL, blue, n = 12) and TIC (red, n = 29) for septal, anterior, and posterior leaflets. The (+) indicates that the anterior leaflet data was previously published and can be found in [17]. Statistical significance between animal group (CTL vs. TIC) is denoted by (**) for p < 0.01 using one-tailed pairwise tests. Statistical significance between leaflets is denoted by (‡) for septal vs. anterior, (#) for anterior vs. posterior, and (§) for septal vs. posterior, all at p ≤ 0.0001, using two-tailed pairwise tests.

### 2.3 Histology and Immunohistochemistry

After thawing, we cut and fixed radial strips (from near-annulus to free edge) of all leaflets in 10% neutral buffered formalin for 24 hours and then transferred to 70% ethanol for long-term storage. The fixed leaflet strips were shipped to a histological service (HistoServ Inc., Amaranth, MD) for embedding, sectioning at 5 µm slice thickness, and staining with Hematoxylin & Eosin and immunohistochemistry. Using a light microscope (Ti2-E High-content Analysis System, Nikon, Tokyo, Japan) with a 10x objective, we acquired full section images for all histological and immunohistochemistry sections. After imaging, we used ImageJ (National Institutes of Health, Bethesda, MD, USA) to measure the total length of the tissue section along the atrial surface and separated each section into three equidistant regions (i.e., near-annulus, belly, and free edge). We then measured the normal vectors between the atrial and ventricular surfaces at three random positions within each region. Finally, we averaged all regional thickness measurements into the total leaflet thickness for each sample.

We assessed four cellular markers associated with remodeling processes: (i) alpha-smooth muscle actin (a-SMA) (Abcam, ab5694, 1:600, Cambridge, MA, US), (ii) Ki67 (Abcam, ab15580, 1:1250, Cambridge, MA, US), (iii) matrix-metalloprotease 13 (MMP13) (Abcam, ab39012, 1:50, Cambridge, MA, US), and (iv) transforming growth factor beta-1 (TGF-/31) (Abcam, ab9758, 1:100, Cambridge, MA, US). A custom MATLAB program segmented the full cross-sectional leaflet image into three equidistant length regions (near-annulus, belly, and free edge). We then interpolated ten thickness regions of the leaflet cross-sectional images between the atrial and ventricular surfaces, resulting in a total of 30 regions. Each region underwent analysis through our validated custom MATLAB program to spatially resolve the presence of positive pixels and normalize their percentage to the total pixels in that region [17].

### 2.4 Biaxial testing

We tested leaflet samples and analyzed the biaxial mechanics using the methods fully described in our prior work [17,21]. First, we isolated a 7x7 mm square sample from the belly region of each septal, anterior, and posterior leaflet. We then applied four fiducial markers to the atrial surface of each sample to enable strain tracking during testing. Before testing, we measured sample thickness at four locations using a digital thickness gauge (547-500S, Mitutoyo Corp., Kawasaki, Japan) and captured an image of the stress-free configuration on a calibrated grid. Next, we mounted all samples on a biaxial testing device (Biotester, CellScale, Waterloo, ON, Canada) submerged in 37°C 1x PBS, and performed ten preconditioning cycles of equibiaxial extension to 300 mN followed by two final equibiaxial cycles to 300 mN. While testing, we continuously captured circumferential and radial forces and images of the fiducial markers at 5 Hz.

We used the image analysis software Labjoy (CellScale, Waterloo, ON, Canada) to track the coordinates of the fiducial markers throughout testing. Our analysis focused on the downstroke of the last loading cycle, and we reported membrane tension-stretch data relative to the preloaded stress-free configuration. To characterize tissue deformation, we computed the deformation gradient tensor F relative to the stress-free reference state and derived the right Cauchy-Green deformation tensor C F^T^F to determine in-plane stretches throughout the experiment. Membrane tension was calculated by dividing the measured force by the perpendicular rake-to-rake distance in the deformed configuration at each time point.

To quantitatively compare the nonlinear membrane tension-stretch curves between CTL and TIC for all three leaflets, we analyzed four characteristic metrics of the resulting nonlinear J-shaped curves: (1) toe stiffness, defined as the slope in the lower linear region of the curve; (2) calf stiffness, measured as the slope near 20 N/m membrane tension; (3) transition stretch, identified as the stretch value at the curve’s inflection point (approximated by the intersection of toe and calf stiffness lines); and (4) anisotropy index, calculated as the ratio of circumferential to radial stretches at 20 N/m tension.

### 2.5 Two-Photon Microscopy

We imaged and analyzed the collagen microstructure and cell nuclei using the same two-photon microscopy methods described in our previous work [21,25]. The mechanically tested square sample from the leaflet belly was briefly stained with Hoechst 33342 (Thermo Fischer Scientific, Hoechst 33342, Waltham, MA, US) for 20 minutes to visualize the cell nuclei. The tissue sample was then optically cleared under sonication for 30 minutes in a solution containing glycerol:DMSO: 5x PBS at 50:30:20% (v/v), respectively [26]. Next, we imaged the leaflet samples with a two-photon microscope (Ultima IV, Bruker, Billerica, MA, USA) to visualize collagen fiber orientation via second harmonic generation (SHG) and cell nuclei via fluorescent excitation, respectively. Before imaging, the leaflet samples were placed on a foil-lined glass slide with a coverslip to flatten the sample during imaging without imposing any stresses. We captured images from the atrial to the ventricular surface at three centrally located regions (500x500 µm, for each region) using a 20x water-immersion objective (Olympus, XLUMPLFLN, Center Valley, PA, US) at an excitation wavelength of 900 nm for SHG and 800 nm for fluorescence. We collected the backscattered SHG signal through a PMT channel filter (460 nm ± 25 nm) and acquired a z-stack of images at a step size of 10 µm. To analyze the SHG images, we used the orientation distribution analysis with the Gaussian gradient method in the ImageJ plugin OrientationJ (National Institutes of Health, Bethesda, MD, USA). We then averaged and interpolated across the three z-stacks for each leaflet sample and then fit a symmetric von Mises distribution to each histogram to estimate the mean fiber angle µ and the fiber orientation concentration K [27]. To analyze the cell nuclei morphology and orientation, we used our custom-validated MATLAB code to quantify individual nuclei contours, nuclear aspect ratio (NAR), and circularity from each image [17]. Similarly, we fit von Mises distributions to the nuclear orientation histograms and normal distributions to the NAR and circularity histograms.

### 2.6 Statistical Analysis

For all morphological, thickness, biaxial, and two-photon distribution data comparisons, we used a linear mixed effects model in R via the package afex (Version 4.4.2). Our models included fixed effects for animal group (CTL vs. TIC), leaflet type (anterior, posterior, septal), direction (X vs. Y) where applicable, and their interactions. Random intercepts were included for subjects to account for repeated measurements within individuals. We performed a Type III ANOVA analysis with Satterthwaite’s method to assess the significance of main effects and interactions. We then performed one-tailed pairwise comparisons between CTL and TIC animals for each leaflet type using the emmeans package in R, and performed two-tailed pairwise comparisons using Tukey’s method to compare between leaflets from the CTL and TIC animals. We defined statistical significance as p-values less than 0.05. We reported data as mean with the standard deviation of the mean.

## 3. RESULTS

### 3.1 Animal model outcomes

We isolated 17 tricuspid valves from CTL sheep and 33 tricuspid valves from TIC sheep. Our study included all tricuspid valve leaflets to assess their geometric and functional changes. Echocardiographic and hemodynamic evaluations revealed significant parameter changes in TIC sheep consistent with biventricular dysfunction and functional TR. Key findings included reduced biventricular ejection fraction, evidence of biventricular remodeling, tricuspid valve annular dilation, and increased severity of TR (Table 1) [17].

### 3.2 TIC-induced changes to leaflet geometry

Leaflet geometry, including area and thickness, is critical for assessing valve function in TR [16]. While prior studies detailed size and shape differences in healthy tricuspid valve leaflets [21], geometric changes in TIC leaflets remain underexplored. Building on our previous findings of geometric changes in TIC anterior leaflets [17], we hypothesized that all TIC leaflets would exhibit distinct geometric changes. To test this, we quantified leaflet area and thickness across all leaflets (Figure 1a, Figures 2a–c).

**Figure 2:**
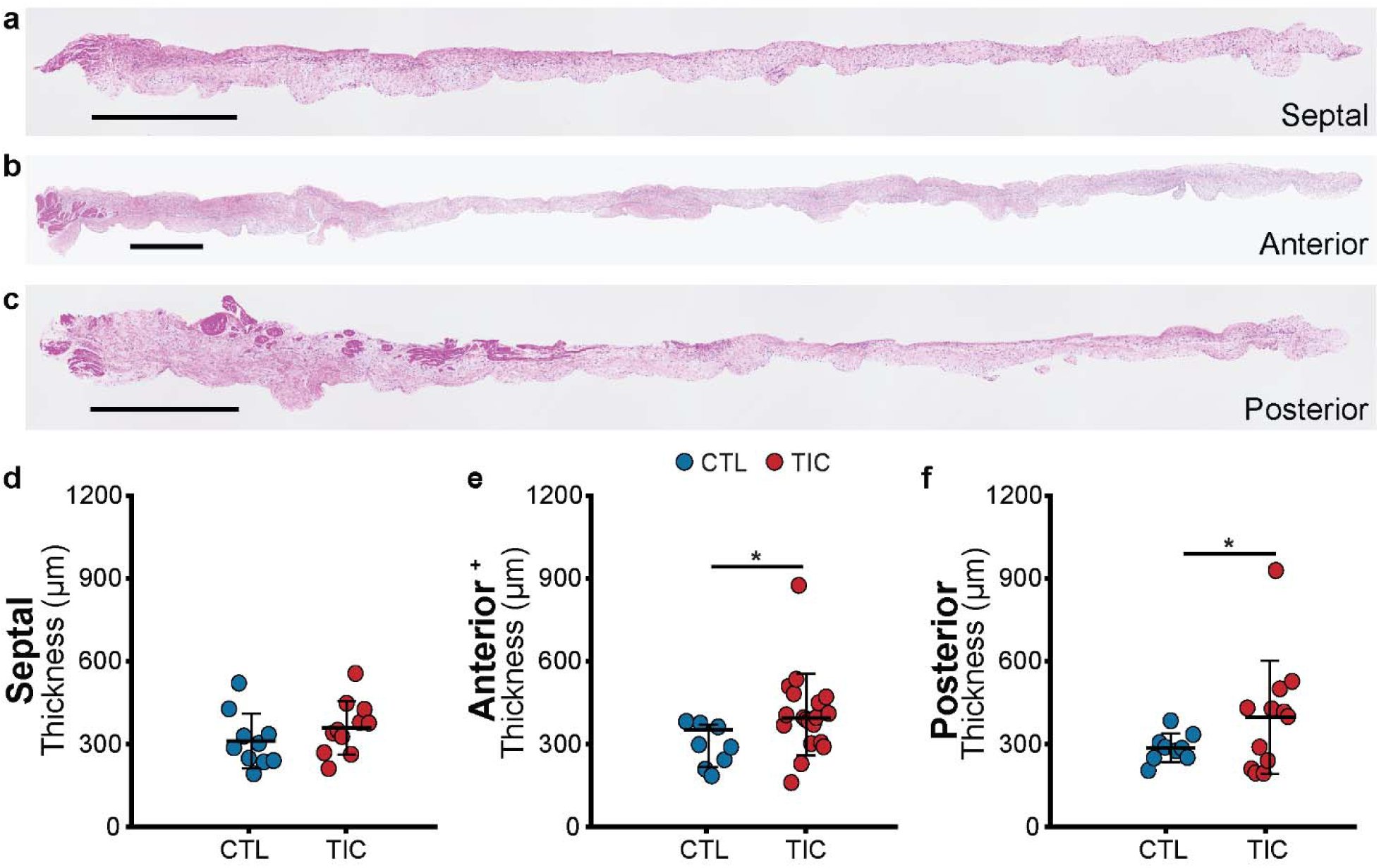
Tricuspid anterior and posterior leaflet thickness increases in sheep with tachycardia-induced cardiomyopathy (TIC). Representative images of (a) septal, (b) anterior, and (c) posterior leaflet strips. Averaged thickness comparisons between control (CTL, blue, n = 8-10) and TIC (red, n = 11-12) (d) septal, (e) anterior, and (f) posterior leaflets. The (+) indicates that the anterior leaflet data was previously published and can be found in [17]. Statistical significance between animal group (CTL vs. TIC) is denoted by (*) for p < 0.05 using one-tailed pairwise tests.

We compared the leaflet area measurements between CTL and TIC animals, and we found that TIC septal and anterior leaflets significantly increased in area compared to respective CTL leaflets (p = 0.009 and p = 0.002, respectively) (Figure 1b-c). In contrast, the TIC posterior leaflets’ area remained unchanged (p = 0.153) (Figure 1d). We found a significant interaction between animal group and leaflet type (p = 0.044), highlighting the heterogeneous remodeling response among leaflets. For instance, in CTL animals, we found that the septal leaflet area was significantly smaller than the anterior and posterior leaflets (both p ≤ 0.0001), while among TIC animals, leaflet area differed significantly across all leaflets (all p ≤ 0.0001). We compared the average thickness between CTL and TIC animals from our histological cross-sectional images, and we found that only the TIC anterior and posterior leaflets significantly increased (p = 0.024 and p = 0.033, respectively) in thickness compared to their respective CTL leaflets (Figures 2d-f). However, we did not detect any significant differences between the leaflets of CTL animals or between the leaflets of TIC animals. These findings highlight the distinct growth of each tricuspid leaflet under disease conditions, suggesting differential remodeling exists between the tricuspid leaflets.

### 3.3 TIC-induced changes to leaflet mechanics

Leaflet remodeling in heart failure patients alters the mechanical properties of the valve leaflets [13]. Building on our previous findings of altered anterior tricuspid leaflet mechanics in TIC animals [17], we hypothesized that each TIC leaflet would exhibit distinct and leaflet-specific changes in mechanical behavior. All leaflets exhibited the classic J-shaped loading curve of biological collagenous tissues (Figures 3a, 3d, 3g) [28]. Our analysis did not detect any significant change in calf stiffnesses between CTL and TIC animals for any of the leaflets, in either the circumferential and radial direction (Figures 3b, 3e, 3h). However, we did find that the TIC anterior leaflet exhibited a significant increase in circumferential toe stiffness when compared to CTL (p = 0.022). In contrast, we failed to detect a significant change in the posterior and septal leaflets’ toe stiffnesses. Our between-leaflet comparison revealed that circumferential toe stiffness differed significantly between the TIC septal and anterior leaflets (p = 0.044) (Figures 3b and 3e).

**Figure 3:**
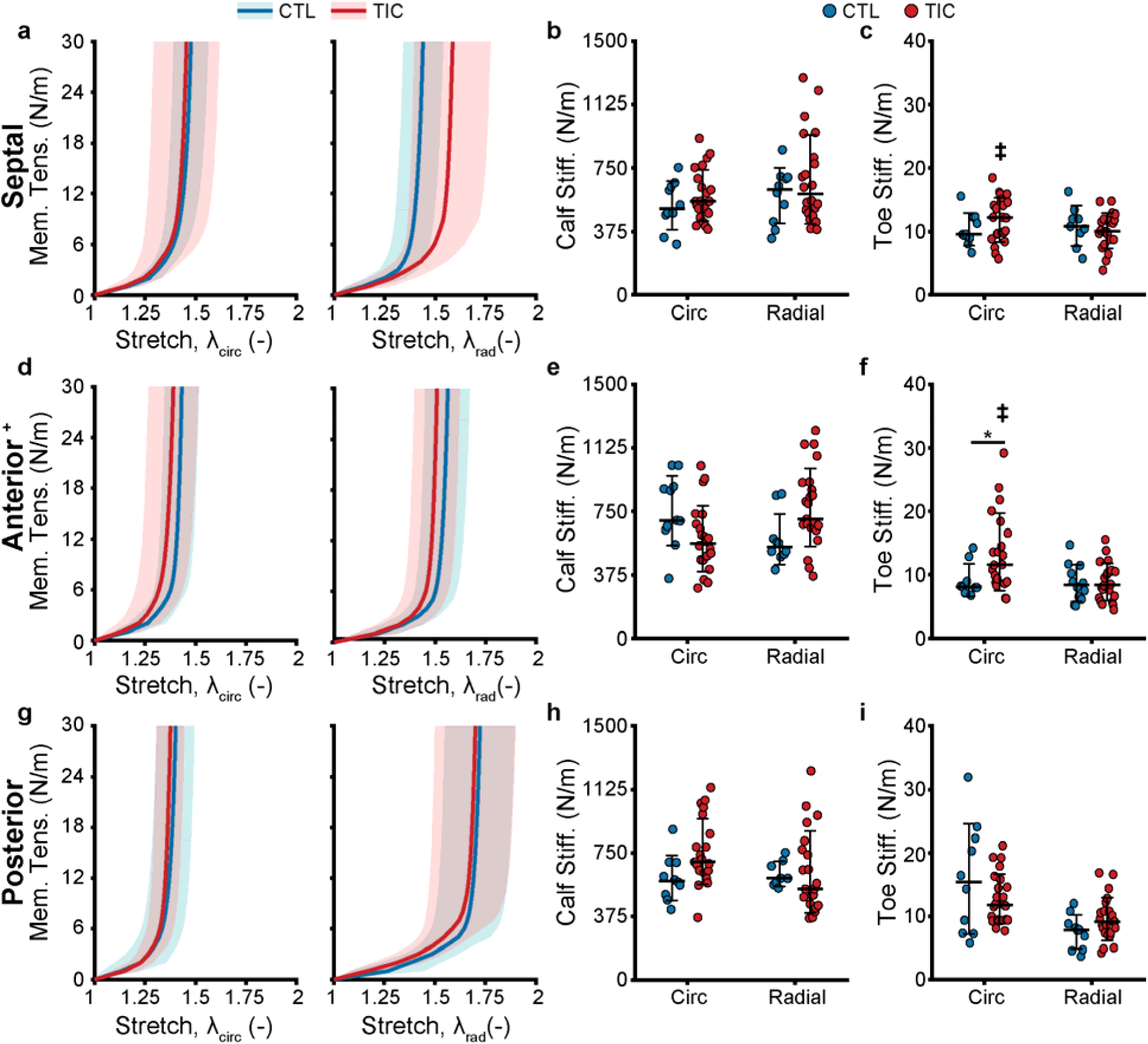
Tachycardia-induced cardiomyopathy (TIC) anterior leaflets are stiffer at low stretches in the circumferential (circ) directions. Control (CTL, blue, n = 10 for each leaflet) and TIC (red, n = 22 for each leaflet) membrane tension (Mem. Tens.) vs stretch average curves (solid) with standard deviation (shaded) in the circumferential and radial (rad) directions for (a) posterior and (d) septal leaflets. Comparisons of the (b & e) stiffness at large stretches (calf stiffness) in the circumferential and radial directions for the posterior and septal leaflets, respectively. Comparisons of the (c & f) stiffness at small stretches (toe stiffness) in the circumferential and radial directions for the posterior and septal leaflets, respectively. The (+) indicates that the anterior leaflet data was previously published and can be found in [17]. Statistical significance between animal group (CTL vs. TIC) is denoted by (*) using one-tailed pairwise tests at p < 0.05, and statistical significance between septal and anterior leaflets is denoted by (‡) using two-tailed pairwise tests at p < 0.05.

The most pronounced between-leaflet differences emerged when comparing the transition stretches and anisotropy indices. We found that the anterior leaflet showed a significant decrease in the circumferential transition stretch between CTL and TIC animals (p = 0.047), while we failed to detect any significant changes in the posterior and septal leaflets (Figure 4 a-c). Our between-leaflet comparison within CTL animals revealed that the posterior leaflet radial transition stretches significantly differed from the other leaflets (anterior-posterior: p = 0.0009; septal-posterior: p ≤ 0.0001). Among the TIC animals, we found that the septal leaflet’s circumferential transition stretch was significantly different from the anterior leaflet (p = 0.036), while the posterior leaflet’s radial transition stretch was significantly different from the other leaflets (anterior-posterior: p ≤ 0.0001; septal-posterior: p = 0.0002). Our analysis of anisotropy indices revealed a near-significant interaction effect between animal group and leaflet type (p = 0.052). We found that the septal leaflet anisotropy index significantly decreased (p = 0.004) between CTL and TIC animals to a value below 1, indicating larger stretches in the radial direction (Figure 4d). In contrast, the anterior and posterior leaflets retained stable anisotropic properties (Figures 4e and 4f). Our between-leaflet comparison within CTL animals revealed significant differences between all leaflets (septal-anterior: p = 0.024; anterior-posterior: p = 0.021; septal-posterior: p ≤ 0.0001). Among TIC animals, the posterior leaflet’s anisotropy differed significantly from both the other leaflets (anterior-posterior: p = 0. 0009; septal-posterior: p = 0.0005), revealing a disease-induced shift in the septal leaflet’s radial behavior. Overall, TIC induced the most significant remodeling mechanical changes in the septal leaflet. While the CTL septal leaflet exhibited greater circumferential compliance, the TIC septal leaflet remodeled in a manner that resulted in greater radial compliance, aligning its behavior more closely with the anterior and posterior leaflets. These findings underscore the pronounced remodeling of the septal leaflet under disease conditions, leading to altered anisotropy and compliance patterns.

**Figure 4:**
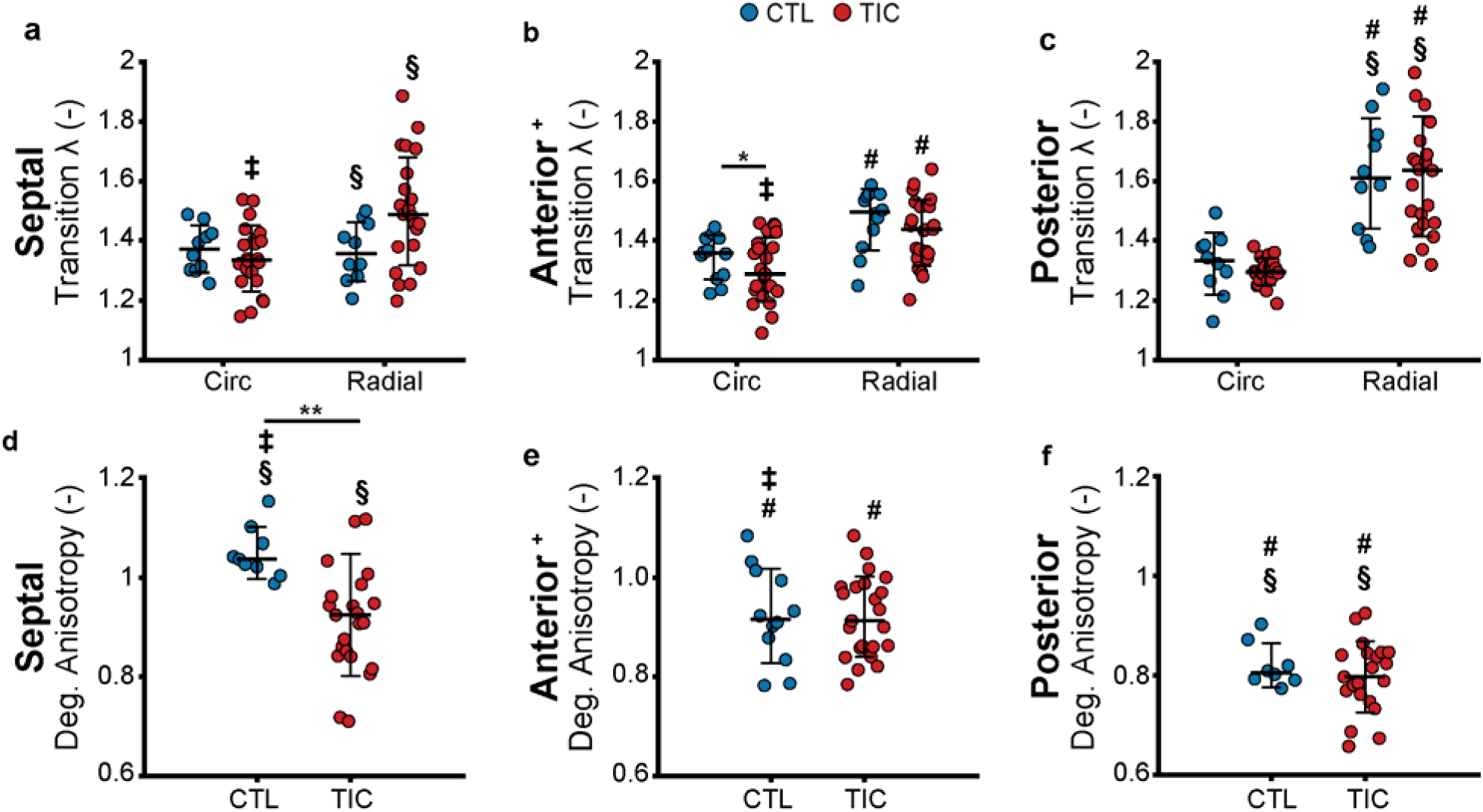
Tachycardia-induced cardiomyopathy (TIC) septal leaflets exhibit increased radial transition stretch and decreased anisotropy. Comparisons of the transition stretch in circumferential and radial directions between control (CTL, blue, n = 10) and TIC (red, n = 22) (a) septal, (b) anterior, and (c) posterior leaflets. Comparisons of the degree of anisotropy between CTL (blue, n = 10) and TIC (red, n = 22) (d) septal, (e) anterior, and (f) posterior leaflets. The (+) indicates that the anterior leaflet data was previously published and can be found in [17]. Statistical significance between animal group (CTL vs. TIC) is denoted by (*) for p < 0.05 and (**) for p < 0.01 using one-tailed pairwise tests. Statistical significance between leaflets is denoted by (‡) for septal vs. anterior, (#) for anterior vs. posterior, and (§) for septal vs. posterior, all at p < 0.05, using two-tailed pairwise tests.

### 3.4 Increased expression of cellular remodeling markers in TIC leaflets

Pathological activation of valvular endothelial cells (VECs) and valvular interstitial cells (VICs) promotes extracellular matrix (ECM) remodeling through expression, secretion, and degradation of load-bearing proteins [29]. We hypothesized that each TIC leaflet would exhibit elevated expression of remodeling markers, with variations in the magnitude and region of expression between leaflets. Specifically, we anticipated increased expression of all markers tested (a-SMA, Ki67, MMP13, TGF-/31) [17,18].

In the TIC septal leaflets, we observed region-specific increases in all four remodeling markers, particularly in the near-annulus and belly regions, while MMP13 expression decreased at the free edge (Figure 5). These findings align with our previous observations of increased remodeling marker expression in TIC anterior leaflets (Figure 6). In the TIC posterior leaflets, expression of a-SMA and Ki67 increased in the near-annulus region (Figures 7a and 7b). In contrast, the expression levels of MMP13 and TGF-/31 decreased throughout the TIC posterior leaflets (Figures 7c and 7d). Additionally, all TIC leaflets exhibited regionally increased cell nuclei density consistent with elevated Ki67 expression, see Supplementary Figure S1.

**Figure 5:**
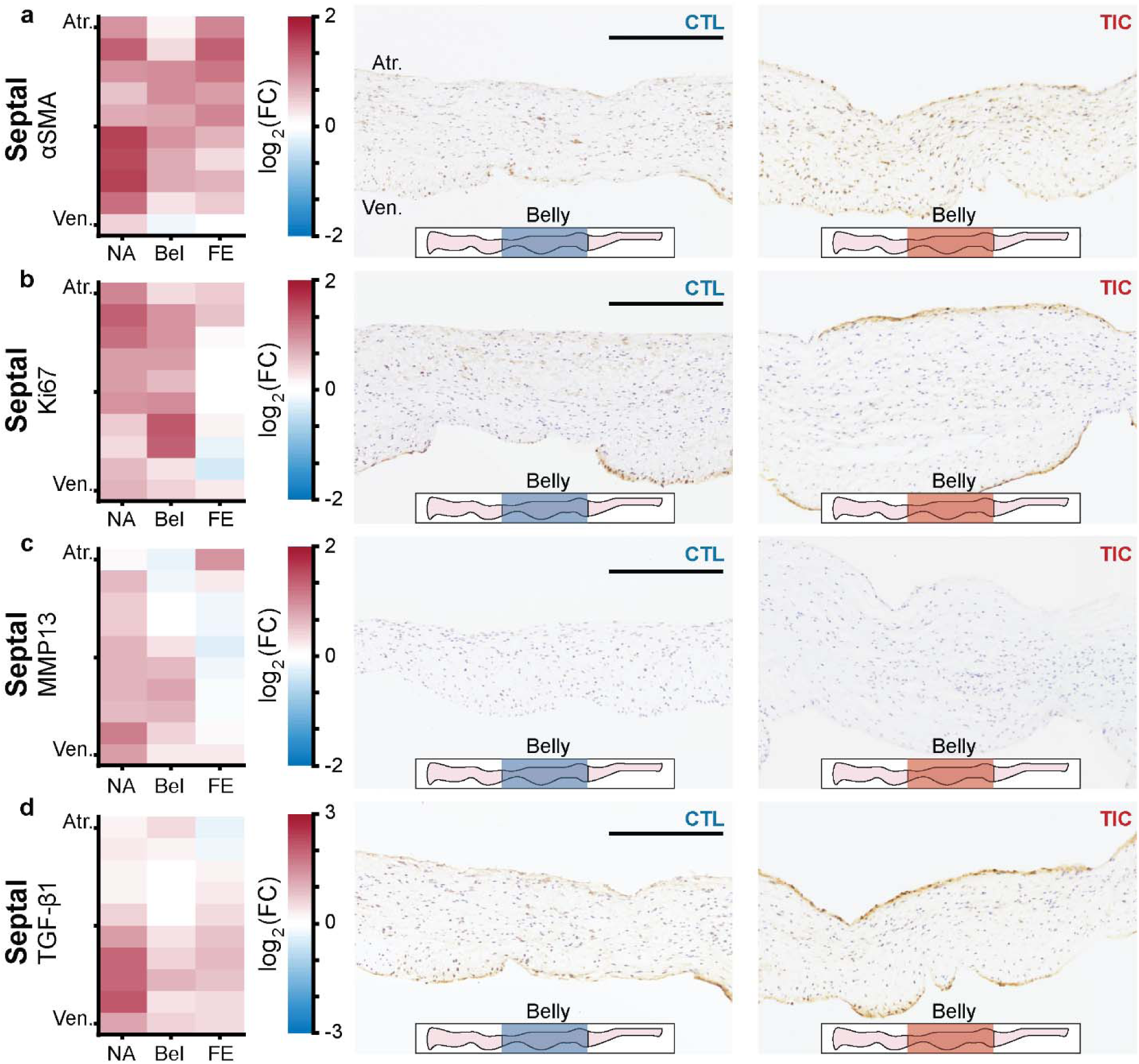
Changes in remodeling cellular markers in tachycardia-induced cardiomyopathy (TIC) septal leaflets. (a-d) Heat maps (left) showing regional expression of (a) alpha-smooth muscle actin (a-SMA), (b) Ki67, (c) matrix metalloproteinase 13 (MMP13), (d) transforming growth factor beta (TGF-/31). The heatmaps are separated into three equidistant regions along the radial axis (near-annulus (NA), belly (Bel), free edge (FE)) and 10 thickness regions from the atrial surface (Atr.) to the ventricular surface (Ven.). Fold change (FC) between control (CTL, n = 6) and TIC (n = 6) was determined by the ratio of positively stained pixel percentage between TIC and CTL. Color map indicates the logarithm base 2 of the fold change, interpreted as (positive, red): TIC expression is higher than CTL, (0, white): TIC and CTL expression are approximately equal, and (negative, blue): TIC expression is less than CTL. Representative images of CTL (middle) and TIC (right) posterior leaflets are shown with the atrial surface upward. The inset leaflet schematic depicts the longitudinal region from which the representative image was taken. Scale bars = 500 µm.

**Figure 6:**
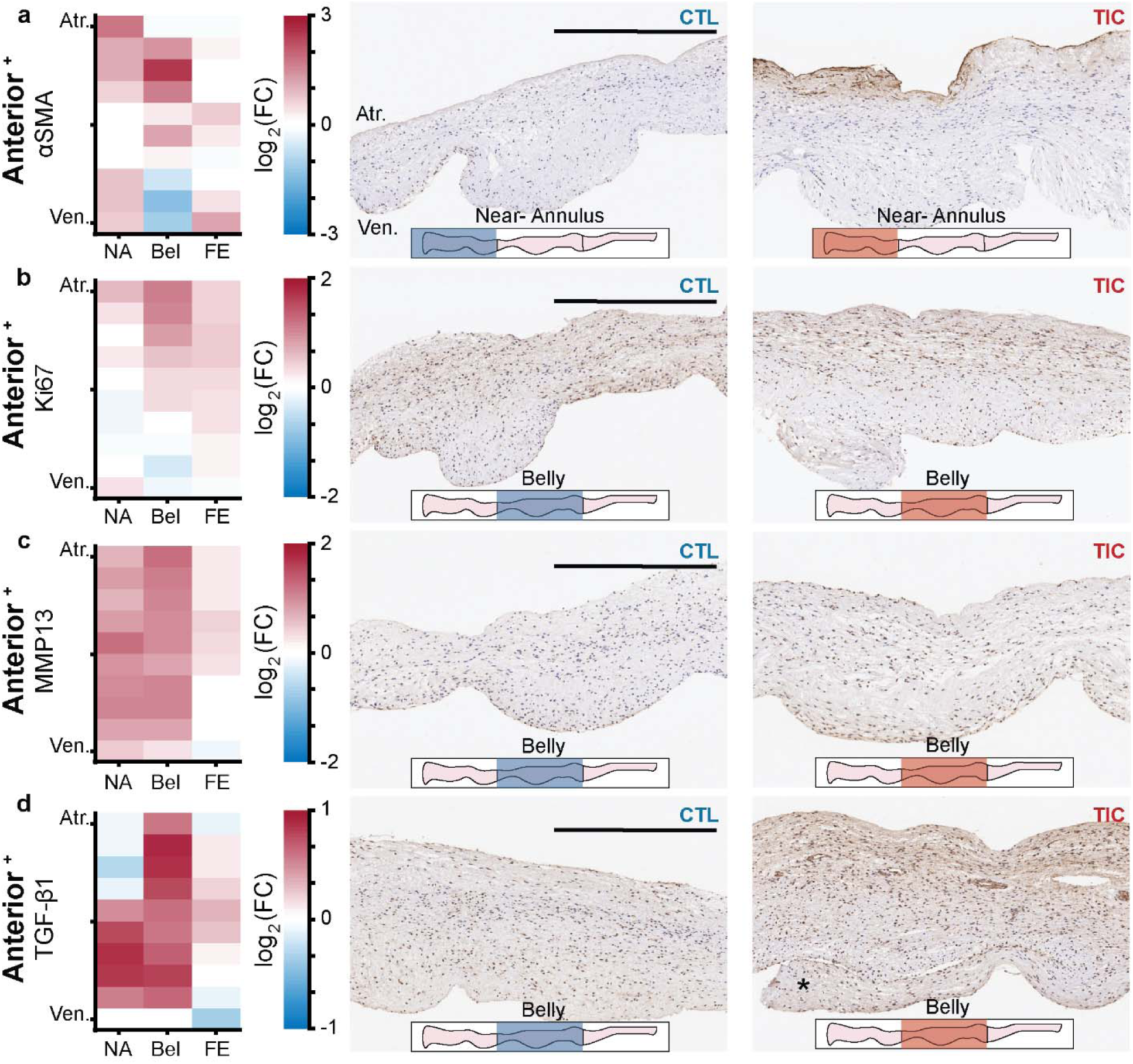
Changes in remodeling cellular markers in tachycardia-induced cardiomyopathy (TIC) anterior leaflets. The (+) indicates that the anterior leaflet data was previously published and can be found in [17]. (a-d) Heat maps (left) showing regional expression of (a) alpha-smooth muscle actin (a-SMA), (b) Ki67, (c) matrix metalloproteinase 13 (MMP13), (d) transforming growth factor beta (TGF-/31). The heatmaps are separated into three equidistant regions along the radial axis (near-annulus (NA), belly (Bel), free edge (FE)) and 10 thickness regions from the atrial surface (Atr.) to the ventricular surface (Ven.). Fold change (FC) between control (CTL, n = 6) and TIC (n = 6) was determined by the ratio of positively stained pixel percentage between TIC and CTL. Color map indicates the logarithm base 2 of the fold change, interpreted as (positive, red): TIC expression is higher than CTL, (0, white): TIC and CTL expression are approximately equal, and (negative, blue): TIC expression is less than CTL. Representative images of CTL (middle) and TIC (right) posterior leaflets are shown with the atrial surface upward. The inset leaflet schematic depicts the longitudinal region from which the representative image was taken. Scale bars = 500 µm.

**Figure 7:**
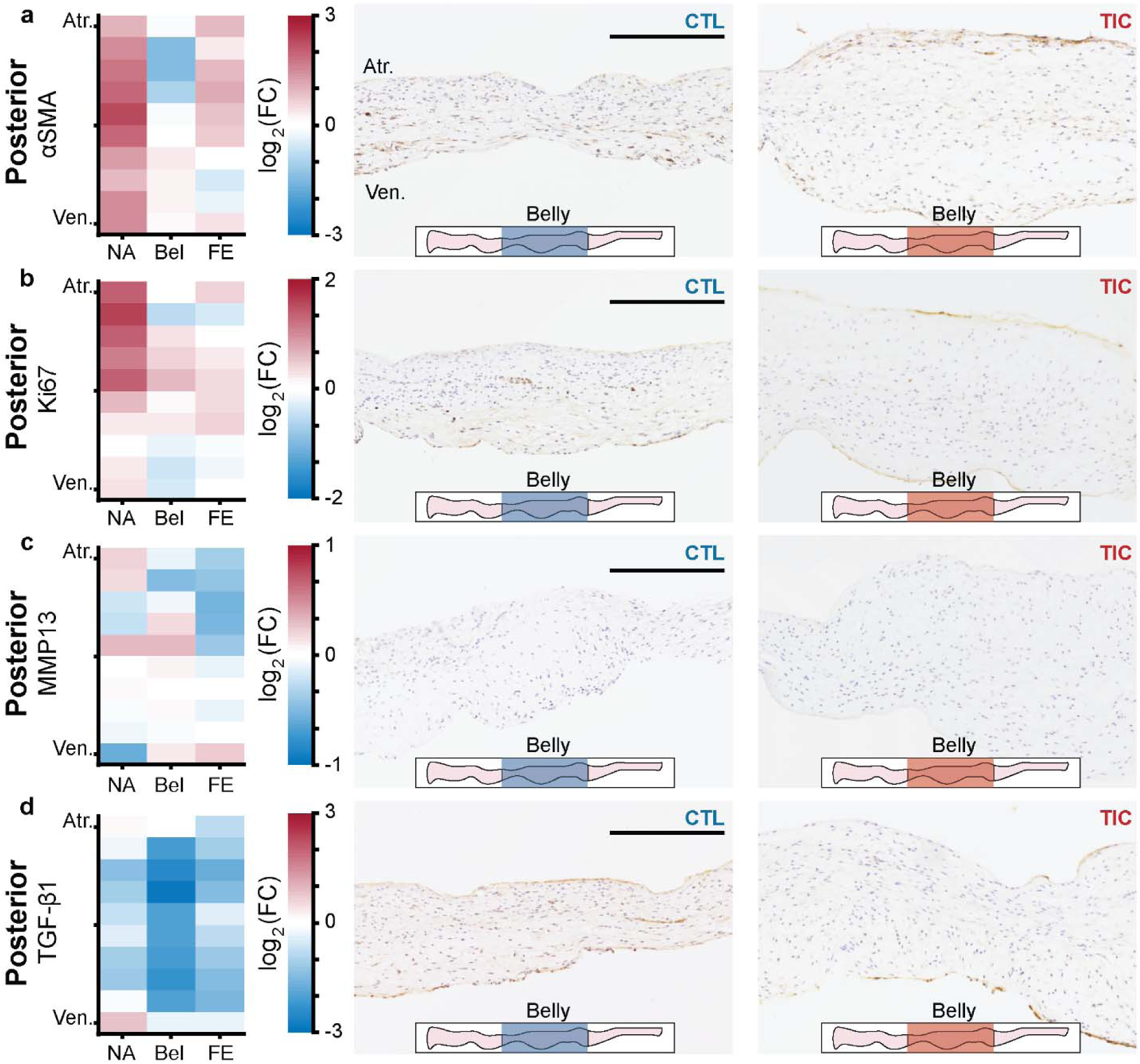
Changes in remodeling cellular markers in tachycardia-induced cardiomyopathy (TIC) posterior leaflets. (a-d) Heat maps (left) showing regional expression of (a) alpha-smooth muscle actin (a-SMA), (b) Ki67, (c) matrix metalloproteinase 13 (MMP13), (d) transforming growth factor beta (TGF-/31). The heatmaps are separated into three equidistant regions along the radial axis (near-annulus (NA), belly (Bel), free edge (FE)) and 10 thickness regions from the atrial surface (Atr.) to the ventricular surface (Ven.). Fold change (FC) between control (CTL, n = 6) and TIC (n = 6) was determined by the ratio of positively stained pixel percentage between TIC and CTL. Color map indicates the logarithm base 2 of the fold change, interpreted as (positive, red): TIC expression is higher than CTL, (0, white): TIC and CTL expression are approximately equal, and (negative, blue): TIC expression is less than CTL. Representative images of CTL (middle) and TIC (right) posterior leaflets are shown with the atrial surface upward. The inset leaflet schematic depicts the longitudinal region from which the representative image was taken. Scale bars = 500 µm.

### 3.5 Changes in collagen fiber orientation in TIC leaflets

Collagen fiber orientation is a critical determinant of leaflet stiffness, ensuring mechanical functionality during the cardiac cycle [30]. Building on previous findings of increased fiber dispersion in the TIC anterior leaflets [17], we hypothesized that all TIC leaflets would exhibit increased collagen fiber dispersion. To test this, we fit von Mises probability distribution functions to collagen fiber orientation histograms for leaflets from CTL and TIC animals (Figures 8a, 8c, 8e). We quantified the concentration parameter K from the collagen fiber orientation histograms and measured fiber dispersion as a function of imaging depth. A large K value indicates more aligned fibers, while lower values suggest greater dispersion and more isotropic distribution of collagen fibers.

**Figure 8:**
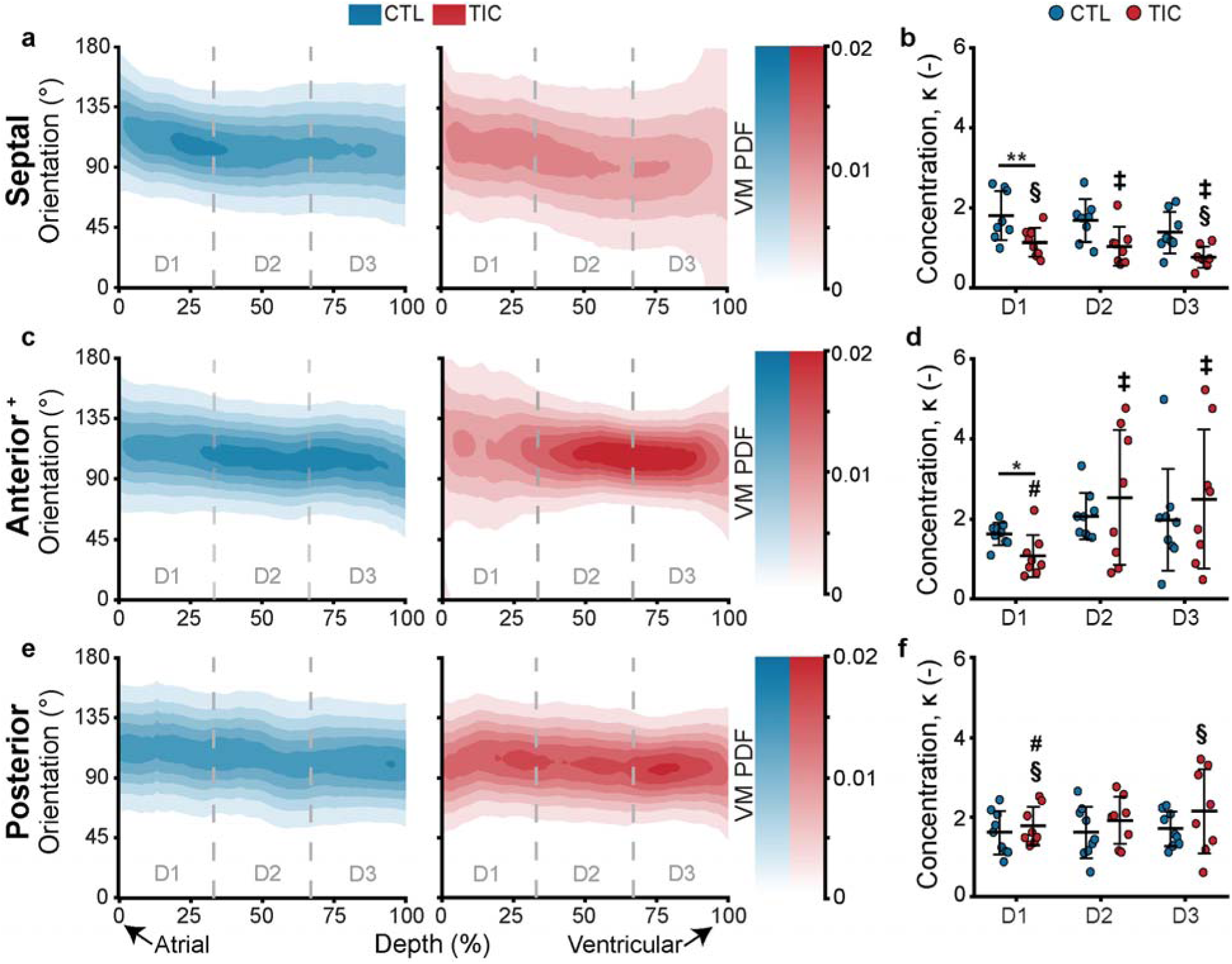
Through-depth collagen microstructure reveals similar mean orientations and regional concentration changed throughout depth between control (CTL) and tachycardia-induced cardiomyopathy (TIC) tricuspid leaflets. Heat map visualizations of von Mises probability distribution functions (VM PDF) fit to collagen fiber orientation histograms of two-photon acquired images throughout the entire depth (0% - Atrial surface, 100% - Ventricular surface) of averaged CTL (blue, n = 9 for each leaflet) and TIC (red, n = 9 for each leaflet) (a) septal, (c) anterior, (e) posterior leaflets. Comparisons of VM concentration parameter, K, across depth regions D1, D2, and D3 for the (b) septal, (d) anterior, and (f) posterior leaflets. The (+) indicates that the anterior leaflet data was previously published and can be found in [17]. Statistical significance between animal group (CTL vs. TIC) is denoted by (*) for p < 0.05 and (**) for p < 0.01 using one-tailed pairwise tests. Statistical significance between leaflets is denoted by (‡) for septal vs. anterior, (#) for anterior vs. posterior, and (§) for septal vs. posterior, all at p < 0.05, using two-tailed pairwise tests.

Our analysis revealed that the TIC septal and anterior leaflet significantly increased in fiber dispersion near the atrial surface (p = 0.005 and p = 0.011, respectively) (Figures 8b and 8d). In contrast, we failed to detect any significant changes in the fiber dispersion in the other depth regions and throughout the posterior leaflet. Our between-leaflet comparison revealed that a significant difference in fiber dispersion was only present in the TIC animals. We found that the TIC posterior leaflet exhibited significantly different K values from the other leaflets in the 0-33% depth (D1), nearest the atrial surface (anterior-posterior: p = 0.017; septal-posterior: p = 0.033). In the 33-67% depth (D2), we found that the TIC septal leaflet was significantly different (p = 0.004) from the anterior leaflet. Additionally, we found that the TIC septal leaflet significantly differed from the other leaflets in the 67-100% depth (D3), nearest the ventricular surface (septal-anterior: p = 0. 0.005; septal-posterior: p = 0.028). Similar analyses revealed cell nuclei of TIC septal leaflets concentrating in the circumferential direction within the 0-33% depth, see Supplementary Fig. 2. In contrast, we observed no notable difference in cell nuclei orientation, nuclear aspect ratio, or circularity between CTL and TIC anterior and posterior leaflets, see Supplementary Fig. 3 and Fig. 4. Overall, we observed that TIC-induced changes resulted in more radially dispersed collagen fibers in TIC septal and anterior leaflets near the atrial surface.

## 4. DISCUSSION

Our study provides a comprehensive analysis of the heterogeneous remodeling responses of tricuspid valve leaflets in a biventricular heart failure model, highlighting leaflet-specific changes at the tissue and cellular levels. The distinct morphological and mechanical properties between the tricuspid leaflets are essential for maintaining healthy valve function [21,31]. Our findings reveal that biventricular heart failure leads to heterogeneous remodeling between the tricuspid leaflets.

At the cellular level, we observed localized profibrotic and degradative responses driving heterogeneous ECM remodeling both between and within tricuspid leaflets [32]. Elevated expression of a-SMA in all TIC leaflets indicates VIC activation as a central mechanism in leaflet remodeling [33,34]. This activation appears localized, with distinctive changes in a-SMA expression within each leaflet, suggesting region-specific remodeling processes. Activated VICs assume a hypersecretory phenotype – known as myofibroblasts – contributing to fibrosis by collagen secretion [33]. The expression of MMP13 is indicative of ECM degradation [17]. In the TIC septal and anterior leaflets’ near-annulus and belly regions, elevated expression of these remodeling markers (a-SMA and MMP13) indicate increased matrix turnover. Conversely, the decreased expression of a-SMA and MMP13 in the free edge region of septal and posterior leaflets signify limited ECM turnover. The localization of both Ki67 and TGF-/31 to the endothelial layer TIC leaflets provides evidence of endothelial-to-mesenchymal transition (EndoMT). These findings highlight the critical role of endothelial cells in responding to mechanical stress and regulating fibrotic responses [35]. This regional variation in remodeling markers highlights the leaflet- and region-specific stimuli, possibly related to mechanical loading or biochemical environment, which drive the localized remodeling response [36]. Spatially heterogeneous pathological strains imposed by annular dilation or leaflet tethering trigger these responses [37]. Such strains, often localized along the free edge, trigger mechanobiological pathways promoting remodeling within that region of the TIC leaflets [29,38].

The tissue and cellular-level changes observed in the tricuspid leaflets after TIC provide mechanistic insights into the functional alterations observed in previous in vivo studies of TIC sheep models [39,40]. The TIC septal leaflet exhibited a complex remodeling response characterized by significant geometric, mechanical, and microstructural changes. We observed an increase in leaflet area and a shift in the anisotropy index towards greater radial compliance. This was accompanied by increased collagen fiber dispersion near the atrial surface and elevated expression of a-SMA and MMP13 in the near-annulus and belly regions. These findings collectively indicate enhanced matrix turnover and suggest a compensatory mechanism to accommodate the increased septal-lateral annular dilation observed in TIC animals [32]. The pronounced remodeling observed in the TIC anterior leaflet aligns with previous findings of increased strain in this leaflet during in vivo studies [39]. We observed a substantial increase in leaflet area and thickness, accompanied by increased circumferential toe stiffness and decreased circumferential transition stretch. These changes indicate a tissue and cellular-level response to maintain coaptation and valve function in the face of increased area strain and annular dilation associated with TIC [17,39]. In contrast, the posterior leaflet displayed a more stable mechanical profile, with no significant changes in stiffness or transition stretch. However, localized increases in a-SMA and Ki67 expression suggest that while the posterior leaflet may not undergo extensive structural remodeling, it remains responsive to pathological stimuli.

The heterogeneous remodeling response observed in our biventricular heart failure model reflects the unique properties of each tricuspid leaflet and the differential mechanical stimuli they experience [20,21]. The TIC model induces complex changes across the tricuspid valve apparatus, including right ventricular remodeling, asymmetric annular dilation, and altered chordal tethering forces [22,39,40]. These extrinsic factors create a diverse mechanical environment, subjecting each leaflet to unique biomechanical stimuli and influencing their individual propensity to remodel.

Incorporating these findings into future computational models can significantly enhance our understanding of valvular disease progression, potentially leading to more clinically relevant results. Computational models, informed by the differential remodeling responses observed in the tricuspid valve leaflets, may help simulate various disease scenarios and predict the impact of different therapeutic interventions [19,41]. This approach may allow for a better understanding of the underlying mechanisms driving leaflet-specific remodeling and aid in developing more precise and effective treatment strategies.

While our study provides valuable insights into tricuspid valve leaflet remodeling in the context of biventricular heart failure, several limitations warrant consideration. The TIC sheep model replicates key features of biventricular heart failure, but induced disease within weeks rather than years [22]. Therefore, the observed leaflet changes likely reflect early adaptive responses. Although the TIC sheep model successfully induced differential leaflet growth and remodeling, we failed to establish clear correlations between these adaptive responses and potential stimuli like TR severity or right ventricular dilation. Spatial variations in local mechanical properties and strain distributions within each leaflet may influence the expression of remodeling-associated genes and lead to heterogeneous maladaptive responses in VICs across different leaflet regions [31,42]. Beyond our regional immunohistochemistry analysis of cellular remodeling markers, our study did not fully explore the potential impact of mechanical microenvironments within individual leaflets. Additionally, interspecies differences in cardiac structure and response to injury limit the direct translation of these findings to the pathogenesis of functional TR in humans. To overcome these limitations, future studies may explore long-term disease progression models and spatially heterogeneous remodeling responses within tricuspid valve leaflets.

## 5. CONCLUSION

In conclusion, our study highlights the differential remodeling responses of tricuspid valve leaflets in biventricular heart failure, emphasizing the unique remodeling changes each tricuspid leaflet undergoes. These findings underscore the importance of considering leaflet-specific properties in understanding tricuspid valve dysfunction. By providing insights into how mechanical and cellular factors contribute to these differences, our work paves the way for more targeted and effective interventions in treating tricuspid valve disease.

## DATA AVAILABILITY

The data for Figures 1-8 and their appendices are publicly available on the Texas Data Repository https://doi.org/10.18738/T8/5EPBVN. The histology and immunohistochemistry leaflet images are publicly available on the Texas Data Repository https://doi.org/10.18738/T8/2TRJGV.

## AUTHOR CONTRIBUTIONS

Colton J. Kostelnik: Investigation, Data curation, Formal analysis, Visualization, Writing - original draft, Writing - review & editing. William D. Meador: Conceptualization, Data curation, Software, Formal analysis, Investigation, Methodology, Writing -- original draft, Writing - review & editing. Chien-Yu Lin: Investigation, Writing - review & editing. Mrudang Mathur: Investigation, Writing - review & editing. Marcin Malinowski: Conceptualization, Investigation, Methodology, Writing - review & editing. Tomasz Jazwiec: Investigation, Methodology. Magda L Piekarska: Writing - review & editing. Boguslaw Gaweda: Writing - review & editing. Zuzanna Malinowska: Formal analysis. Tomasz A. Timek: Conceptualization, Resources, Supervision, Funding acquisition, Writing - review & editing. Manuel K. Rausch: Conceptualization, Writing - original draft, Writing - review & editing, Supervision, Funding acquisition.

## ACKNOWLEDGEMENTS

The research reported in this publication was partially supported by the National Heart, Lung, And Blood Institute of the National Institutes of Health under Award Numbers 1R01HL165251-01 and R21HL161832 (MKR), F31HL145976 (WDM), the American Heart Association for their support under Award Number 18CDA34120028 (MKR), and internal grants from Meijer Heart and Vascular Institute at Corewell Health. The opinions, findings, and conclusions expressed are those of the authors and do not necessarily reflect the official views of the National Institutes of Health or the American Heart Association.

The anterior leaflet data reported in this publication was previously published in [17]. This publication expands upon our earlier work by incorporating additional leaflet comparisons to provide new insights into leaflet-specific remodeling as a response to tachycardia-induced cardiomyopathy.

## REFERENCES

[1] J.P. Singh, J.C. Evans, D. Levy, M.G. Larson, L.A. Freed, D.L. Fuller, B. Lehman, E.J. Benjamin, Prevalence and clinical determinants of mitral, tricuspid, and aortic regurgitation (the Framingham Heart Study), The American Journal of Cardiology 83 (1999) 897–902. 10.1016/S0002-9149(98)01064-9.

[2] M. Enriquez-Sarano, D. Messika-Zeitoun, Y. Topilsky, C. Tribouilloy, G. Benfari, H. Michelena, Tricuspid regurgitation is a public health crisis, Progress in Cardiovascular Diseases 62 (2019) 447–451. 10.1016/j.pcad.2019.10.009.

[3] F. Condello, M. Gitto, G.G. Stefanini, Etiology, epidemiology, pathophysiology and management of tricuspid regurgitation: an overview, Reviews in Cardiovascular Medicine 22 (2021) 1115. 10.31083/j.rcm2204122.

[4] E.A. Fender, C.J. Zack, R.A. Nishimura, Isolated tricuspid regurgitation: outcomes and therapeutic interventions, Heart 104 (2018) 798–806. 10.1136/heartjnl-2017-311586.

[5] R.J. Henning, Tricuspid valve regurgitation: current diagnosis and treatment, Am J Cardiovasc Dis. 12 (2022) 1–18.

[6] C.-H. Lee, D.W. Laurence, C.J. Ross, K.E. Kramer, A.R. Babu, E.L. Johnson, M.-C. Hsu, A. Aggarwal, A. Mir, H.M. Burkhart, R.A. Towner, R. Baumwart, Y. Wu, Mechanics of the Tricuspid Valve—From Clinical Diagnosis/Treatment, In-Vivo and In-Vitro Investigations, to Patient-Specific Biomechanical Modeling, Bioengineering 6 (2019) 47. 10.3390/bioengineering6020047.

[7] M. Taramasso, H. Vanermen, F. Maisano, A. Guidotti, G. La Canna, O. Alfieri, The Growing Clinical Importance of Secondary Tricuspid Regurgitation, Journal of the American College of Cardiology 59 (2012) 703–710. 10.1016/j.jacc.2011.09.069.

[8] J.H. Rogers, S.F. Bolling, The Tricuspid Valve: Current Perspective and Evolving Management of Tricuspid Regurgitation, Circulation 119 (2009) 2718–2725. 10.1161/CIRCULATIONAHA.108.842773.

[9] M. Pagnesi, C. Montalto, A. Mangieri, E. Agricola, R. Puri, M. Chiarito, M.B. Ancona, D. Regazzoli, L. Testa, M. De Bonis, N.E. Moat, J. Rodés-Cabau, A. Colombo, A. Latib, Tricuspid annuloplasty versus a conservative approach in patients with functional tricuspid regurgitation undergoing left-sided heart valve surgery: A study-level meta-analysis, International Journal of Cardiology 240 (2017) 138–144. 10.1016/j.ijcard.2017.05.014.

[10] K.J. Grande-Allen, A.G. Borowski, R.W. Troughton, P.L. Houghtaling, N.R. DiPaola, C.S. Moravec, I. Vesely, B.P. Griffin, Apparently normal mitral valves in patients with heart failure demonstrate biochemical and structural derangements, Journal of the American College of Cardiology 45 (2005) 54–61. 10.1016/j.jacc.2004.06.079.

[11] M. Chaput, M.D. Handschumacher, F. Tournoux, L. Hua, J.L. Guerrero, G.J. Vlahakes, R.A. Levine, Mitral Leaflet Adaptation to Ventricular Remodeling: Occurrence and Adequacy in Patients With Functional Mitral Regurgitation, Circulation 118 (2008) 845–852. 10.1161/CIRCULATIONAHA.107.749440.

[12] A. Itoh, G. Krishnamurthy, J.C. Swanson, D.B. Ennis, W. Bothe, E. Kuhl, M. Karlsson, L.R. Davis, D.C. Miller, N.B. Ingels, Active stiffening of mitral valve leaflets in the beating heart, American Journal of Physiology-Heart and Circulatory Physiology 296 (2009) H1766– H1773. 10.1152/ajpheart.00120.2009.

[13] K.J. Grande-Allen, J.E. Barber, K.M. Klatka, P.L. Houghtaling, I. Vesely, C.S. Moravec, P.M. McCarthy, Mitral valve stiffening in end-stage heart failure: Evidence of an organic contribution to functional mitral regurgitation, The Journal of Thoracic and Cardiovascular Surgery 130 (2005) 783–790. 10.1016/j.jtcvs.2005.04.019.

[14] M.K. Rausch, F.A. Tibayan, D. Craig Miller, E. Kuhl, Evidence of adaptive mitral leaflet growth, Journal of the Mechanical Behavior of Biomedical Materials 15 (2012) 208–217. 10.1016/j.jmbbm.2012.07.001.

[15] T.A. Timek, D.T. Lai, P. Dagum, D. Liang, G.T. Daughters, N.B. Ingels, D.C. Miller, Mitral Leaflet Remodeling in Dilated Cardiomyopathy, Circulation 114 (2006). 10.1161/CIRCULATIONAHA.105.000554.

[16] J. Afilalo, J. Grapsa, P. Nihoyannopoulos, J. Beaudoin, J.S.R. Gibbs, R.N. Channick, D. Langleben, L.G. Rudski, L. Hua, M.D. Handschumacher, M.H. Picard, R.A. Levine, Leaflet Area as a Determinant of Tricuspid Regurgitation Severity in Patients With Pulmonary Hypertension, Circ: Cardiovascular Imaging 8 (2015) e002714. 10.1161/CIRCIMAGING.114.002714.

[17] W.D. Meador, M. Mathur, G.P. Sugerman, M. Malinowski, T. Jazwiec, X. Wang, C.M. Lacerda, T.A. Timek, M.K. Rausch, The tricuspid valve also maladapts as shown in sheep with biventricular heart failure, eLife 9 (2020) e63855. 10.7554/eLife.63855.

[18] A. Iwasieczko, M. Gaddam, B. Gaweda, A. Goodyke, M. Mathur, C.-Y. Lin, J. Zagorski, M. Solarewicz, S. Cohle, M. Rausch, T.A. Timek, Valvular complex and tissue remodelling in ovine functional tricuspid regurgitation, European Journal of Cardio-Thoracic Surgery 63 (2023) ezad115. 10.1093/ejcts/ezad115.

[19] M. Mathur, M. Malinowski, T. Jazwiec, T.A. Timek, M.K. Rausch, Leaflet remodeling reduces tricuspid valve function in a computational model, Journal of the Mechanical Behavior of Biomedical Materials 152 (2024) 106453. 10.1016/j.jmbbm.2024.106453.

[20] M. Mathur, T. Jazwiec, W.D. Meador, M. Malinowski, M. Goehler, H. Ferguson, T.A. Timek, M.K. Rausch, Tricuspid valve leaflet strains in the beating ovine heart, Biomech Model Mechanobiol 18 (2019) 1351–1361. 10.1007/s10237-019-01148-y.

[21] W.D. Meador, M. Mathur, G.P. Sugerman, T. Jazwiec, M. Malinowski, M.R. Bersi, T.A. Timek, M.K. Rausch, A detailed mechanical and microstructural analysis of ovine tricuspid valve leaflets, Acta Biomaterialia 102 (2020) 100–113. 10.1016/j.actbio.2019.11.039.

[22] M. Malinowski, A.G. Proudfoot, D. Langholz, L. Eberhart, M. Brown, H. Schubert, J. Wodarek, T.A. Timek, Large animal model of functional tricuspid regurgitation in pacing induced end-stage heart failure, Interactive CardioVascular and Thoracic Surgery 24 (2017) 905–910. 10.1093/icvts/ivx012.

[23] C.J. Kostelnik, K.J. Crouse, J.D. Goldsmith, J.F. Eberth, Impact of cryopreservation on elastomuscular artery mechanics, Journal of the Mechanical Behavior of Biomedical Materials 154 (2024) 106503. 10.1016/j.jmbbm.2024.106503.

[24] G.A. Duginski, C.J. Ross, D.W. Laurence, C.H. Johns, C.-H. Lee, An investigation of the effect of freezing storage on the biaxial mechanical properties of excised porcine tricuspid valve anterior leaflets, Journal of the Mechanical Behavior of Biomedical Materials 101 (2020) 103438. 10.1016/j.jmbbm.2019.103438.

[25] K.J. Smith, M. Mathur, W.D. Meador, B. Phillips-Garcia, G.P. Sugerman, A.K. Menta, T. Jazwiec, M. Malinowski, T.A. Timek, M.K. Rausch, Tricuspid Chordae Tendineae Mechanics: Insertion Site, Leaflet, and Size-Specific Analysis and Constitutive Modelling, Exp Mech 61 (2021) 19–29. 10.1007/s11340-020-00594-5.

[26] W.D. Meador, J. Zhou, M. Malinowski, T. Jazwiec, S. Calve, T.A. Timek, M.K. Rausch, The effects of a simple optical clearing protocol on the mechanics of collagenous soft tissue, Journal of Biomechanics 122 (2021) 110413. 10.1016/j.jbiomech.2021.110413.

[27] A.J. Schriefl, A.J. Reinisch, S. Sankaran, D.M. Pierce, G.A. Holzapfel, Quantitative assessment of collagen fibre orientations from two-dimensional images of soft biological tissues, J. R. Soc. Interface. 9 (2012) 3081–3093. 10.1098/rsif.2012.0339.

[28] M.S. Sacks, Biaxial Mechanical Evaluation of Planar Biological Materials, in: S.C. Cowin, J.D. Humphrey (Eds.), Cardiovascular Soft Tissue Mechanics, Kluwer Academic Publishers, Dordrecht, 2004: pp. 199–246. 10.1007/0-306-48389-0_7.

[29] K.M. Kodigepalli, K. Thatcher, T. West, D.P. Howsmon, F.J. Schoen, M.S. Sacks, C.K. Breuer, J. Lincoln, Biology and Biomechanics of the Heart Valve Extracellular Matrix, JCDD 7 (2020) 57. 10.3390/jcdd7040057.

[30] L.T. Hudson, S.V. Jett, K.E. Kramer, D.W. Laurence, C.J. Ross, R.A. Towner, R. Baumwart, K.M. Lim, A. Mir, H.M. Burkhart, Y. Wu, C.-H. Lee, A Pilot Study on Linking Tissue Mechanics with Load-Dependent Collagen Microstructures in Porcine Tricuspid Valve Leaflets, Bioengineering 7 (2020) 60. 10.3390/bioengineering7020060.

[31] D. Laurence, C. Ross, S. Jett, C. Johns, A. Echols, R. Baumwart, R. Towner, J. Liao, P. Bajona, Y. Wu, C.-H. Lee, An investigation of regional variations in the biaxial mechanical properties and stress relaxation behaviors of porcine atrioventricular heart valve leaflets, Journal of Biomechanics 83 (2019) 16–27. 10.1016/j.jbiomech.2018.11.015.

[32] T.L. Blevins, S.B. Peterson, E.L. Lee, A.M. Bailey, J.D. Frederick, T.N. Huynh, V. Gupta, K.J. Grande-Allen, Mitral Valvular Interstitial Cells Demonstrate Regional, Adhesional, and Synthetic Heterogeneity, Cells Tissues Organs 187 (2008) 113–122. 10.1159/000108582.

[33] H. Ma, A.R. Killaars, F.W. DelRio, C. Yang, K.S. Anseth, Myofibroblastic activation of valvular interstitial cells is modulated by spatial variations in matrix elasticity and its organization, Biomaterials 131 (2017) 131–144. 10.1016/j.biomaterials.2017.03.040.

[34] G.A. Walker, K.S. Masters, D.N. Shah, K.S. Anseth, L.A. Leinwand, Valvular Myofibroblast Activation by Transforming Growth Factor-β: Implications for Pathological Extracellular Matrix Remodeling in Heart Valve Disease, Circulation Research 95 (2004) 253–260. 10.1161/01.RES.0000136520.07995.aa.

[35] Y. Li, K.O. Lui, B. Zhou, Reassessing endothelial-to-mesenchymal transition in cardiovascular diseases, Nat Rev Cardiol 15 (2018) 445–456. 10.1038/s41569-018-0023-y.

[36] K.J. Grande-Allen, J. Liao, The Heterogeneous Biomechanics and Mechanobiology of the Mitral Valve: Implications for Tissue Engineering, Curr Cardiol Rep 13 (2011) 113–120. 10.1007/s11886-010-0161-2.

[37] J.P. Dal-Bianco, E. Aikawa, J. Bischoff, J.L. Guerrero, M.D. Handschumacher, S. Sullivan, B. Johnson, J.S. Titus, Y. Iwamoto, J. Wylie-Sears, R.A. Levine, A. Carpentier, Active Adaptation of the Tethered Mitral Valve: Insights Into a Compensatory Mechanism for Functional Mitral Regurgitation, Circulation 120 (2009) 334–342. 10.1161/CIRCULATIONAHA.108.846782.

[38] S. Korossis, Structure-Function Relationship of Heart Valves in Health and Disease, in: K. Kirali (Ed.), Structural Insufficiency Anomalies in Cardiac Valves, InTech, 2018. 10.5772/intechopen.78280.

[39] T. Jazwiec, M.J. Malinowski, H. Ferguson, J. Parker, M. Mathur, M.K. Rausch, T.A. Timek, Tricuspid Valve Anterior Leaflet Strains in Ovine Functional Tricuspid Regurgitation, Seminars in Thoracic and Cardiovascular Surgery 33 (2021) 356–364. 10.1053/j.semtcvs.2020.09.012.

[40] A. Iwasieczko, T. Jazwiec, M. Gaddam, B. Gaweda, M. Piekarska, M. Solarewicz, M.K. Rausch, T.A. Timek, Septal annular dilation in chronic ovine functional tricuspid regurgitation, The Journal of Thoracic and Cardiovascular Surgery 166 (2023) e393–e403. 10.1016/j.jtcvs.2023.04.003.

[41] C.E. Haese, M. Mathur, C.-Y. Lin, M. Malinowski, T.A. Timek, M.K. Rausch, Impact of tricuspid annuloplasty device shape and size on valve mechanics—a computational study, JTCVS Open 17 (2024) 111–120. 10.1016/j.xjon.2023.11.002.

[42] K.E. Kramer, C.J. Ross, D.W. Laurence, A.R. Babu, Y. Wu, R.A. Towner, A. Mir, H.M. Burkhart, G.A. Holzapfel, C.-H. Lee, An investigation of layer-specific tissue biomechanics of porcine atrioventricular valve anterior leaflets, Acta Biomaterialia 96 (2019) 368–384. 10.1016/j.actbio.2019.06.049.

