## Supplemental Figure 1-3 for "Tricuspid valve leaflet remodeling in sheep with biventricular heart failure: A comparison between leaflets"

**Supplementary Materials**

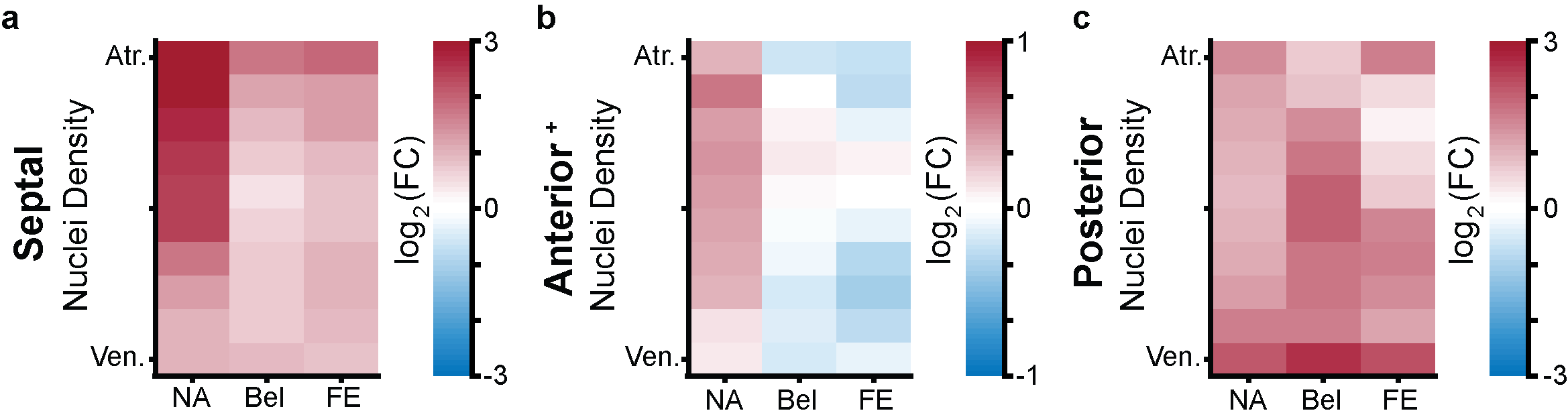

***Figure S1:*** *Heatmaps showing regional cell nuclei density between hematoxylin-eosin stained control (CTL) and tachycardia-induced cardiomyopathy (TIC) (a) septal, (b) anterior, and (c) posterior leaﬂets. The heatmaps are separated into three equidistant longitudinal regions along the leaﬂet’s radial axis (near-annulus) (NA), belly (Bel), and free edge (FE) and ten thickness regions from the atrial surface (Atr.) to the ventricular surface (Ven.) Fold change (FC) between CTL (n = 6) and TIC (n = 6) was determined by the ratio of cell nuclei count between TIC and CTL leaflets. Color map indicates the logarithm base 2 of the fold change, interpreted as (positive, red): TIC leaflet nuclei density is higher than CTL, (0, white): TIC and CTL leaflet nuclei density are approximately equal, and (negative, blue): TIC leaflet nuclei density is less than CTL.*

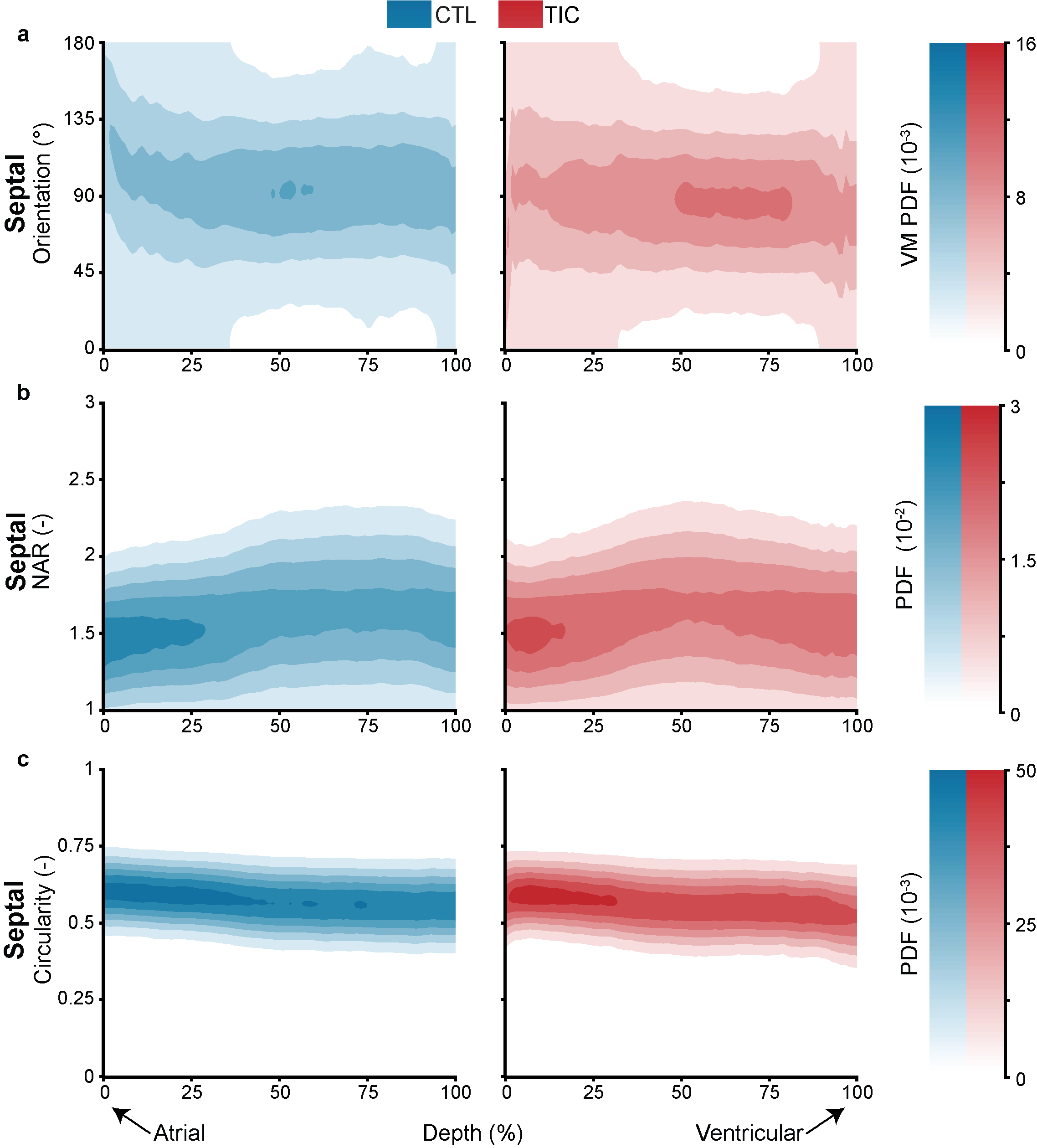

***Figure S2:*** *Through-depth imaging of nuclei orientation, nuclear aspect ratio (NAR), and circularity of control (CTL, blue, n = 9) and tachycardia-induced cardiomyopathy (TIC, red, n = 9) septal leaﬂets.*

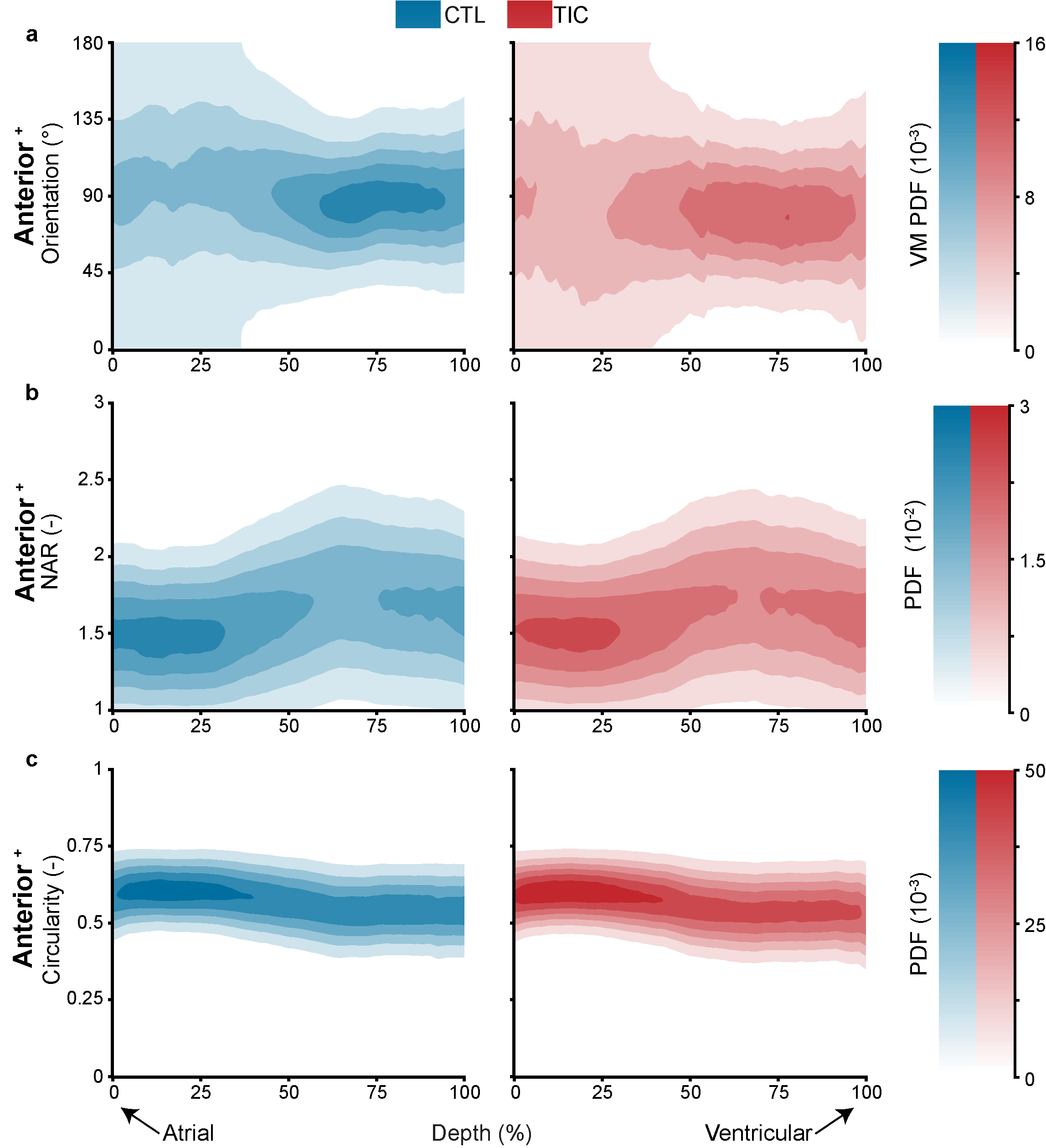

***Figure S3:*** *Through-depth imaging of nuclei orientation, nuclear aspect ratio (NAR), and circularity of control (CTL, blue, n = 9) and tachycardia-induced cardiomyopathy (TIC, red, n = 9) anterior leaﬂets.*

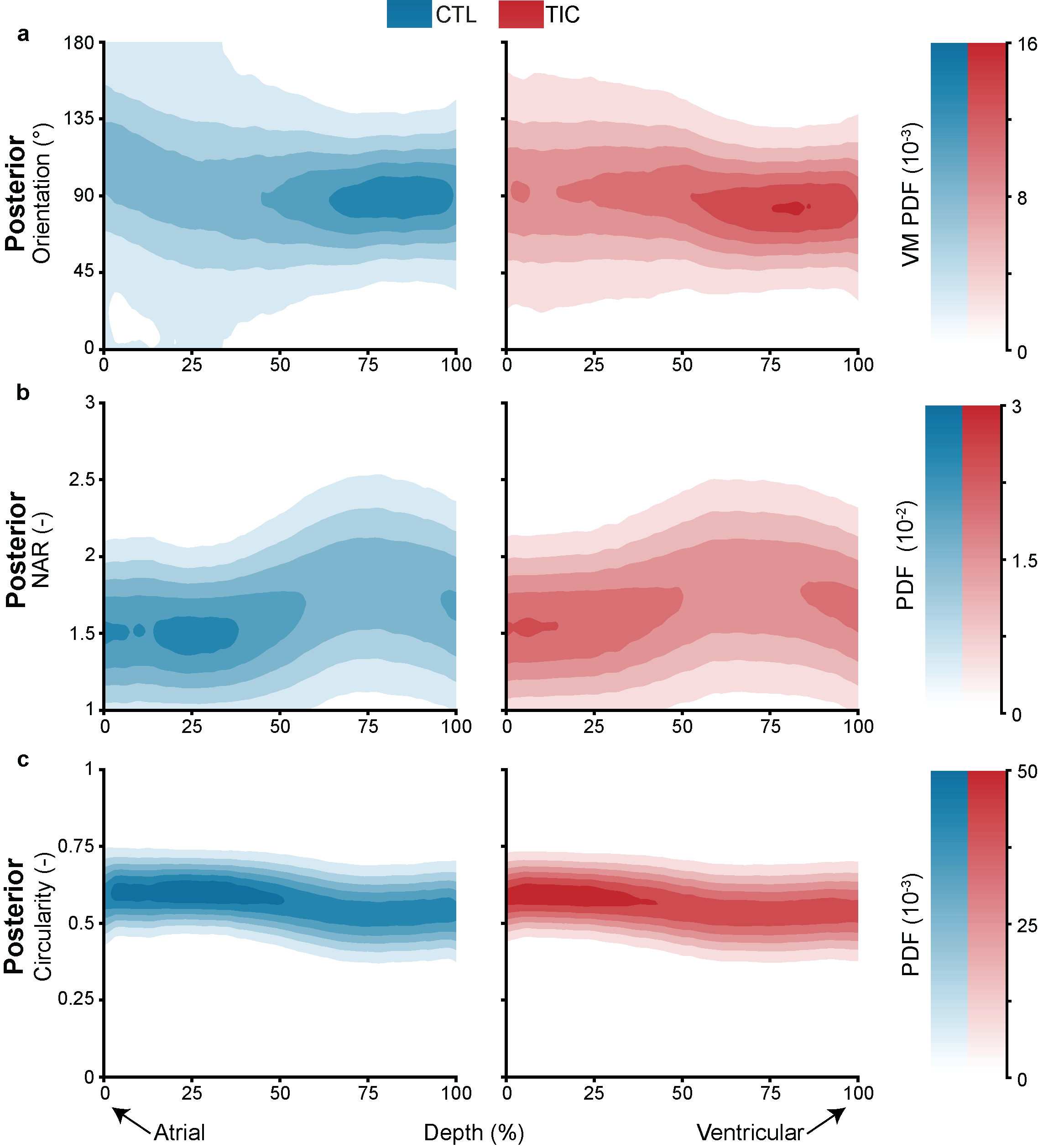

***Figure S4:*** *Through-depth imaging of nuclei orientation, nuclear aspect ratio (NAR), and circularity of control (CTL, blue, n = 9) and tachycardia-induced cardiomyopathy (TIC, red, n = 9) posterior leaﬂets.*
